# Highly accurate prediction of broad-spectrum antiviral compounds with DeepAVC

**DOI:** 10.1101/2025.02.15.638408

**Authors:** Boming Kang, Yang Zhao, Xingyu Chen, Rui Fan, Fuping You, Qinghua Cui

**Affiliations:** Department of Biomedical Informatics, State Key Laboratory of Vascular Homeostasis and Remodeling, School of Basic Medical Sciences, Peking University, 38 Xueyuan Rd, Beijing, 100191, China; Institute of Systems Biomedicine, Beijing Key Laboratory of Tumor Systems Biology, Department of Microbiology & Infectious Disease Center, NHC Key Laboratory of Medical Immunology, Peking University Health Science Center, Peking University, Beijing, China; School of Sports Medicine, Wuhan Sports University, No. 461 Luoyu Rd. Wuchang District, Wuhan 430079, Hubei Province, China

## Abstract

Lethal viruses, such as HIV, pose a significant threat to human life, with each pandemic causing millions of fatalities globally. Small-molecule antiviral drugs provide an efficient and convenient approach to antiviral therapy by either inhibiting viral activity or activating the host immune system. However, conventional antiviral drug discovery is often labor-intensive and time-consuming due to the vast chemical space. Although some existing computational models mitigate this problem, there remains a lack of rapid and accurate method specifically designed for antiviral drug discovery. Here, we propose DeepAVC, a universal framework based on pre-trained large language models, for highly accurate broad-spectrum antiviral compounds discovery, including two models, DeepPAVC for phenotype-based prediction and DeepTAVC for target-based prediction. As a result, DeepAVC greatly outperforms other *in silico* methods. More importantly, in the top predictions, MNS and NVP-BVU972 were identified as novel compounds with promising broad-spectrum antiviral activities by *in vitro* experiments. Finally, DeepAVC demonstrates high interpretability, one of the bottlenecks of current AI methods, due to its ability of analyzing key functional groups of antiviral compounds and important binding sites on targets.

## Introduction

The pandemics caused by deadly infectious viruses, such as Human Immunodeficiency Virus (HIV), Hepatitis C Virus (HCV), Influenza Virus, and Ebola Virus, have posed severe threats to global public health and resulted in substantial economic losses^**1,2**^. Therefore, developing effective antiviral therapies is an urgent priority to address pandemics and save people’s lives. Currently, antiviral therapies can be broadly categorized into small-molecule antiviral drugs and vaccines^**3,4**^. While vaccines are effective at neutralizing viral toxicity, oral antiviral drugs offer flexible dosing and lower production costs, highlighting the importance of small-molecule antivirals^**5,6**^. However, conventional drug screening, design, and optimization are often labor-intensive and time-consuming, with the average development cost has risen to more than $2 billion over a 10-15-year timeline^**7**^. Although some computer-aided drug design strategies, such as molecular docking and quantitative structure-activity relationship modeling, can partially accelerate drug development^**8,9**^, these methods are limited by reliance on biased human prior knowledge and constrained computing speed^**10**^, making it difficult to scale to the vast chemical space of drug-like small-molecule compounds, estimated to be between 10^23^ and 10^60 **11**^.

Recently, advances in machine learning and deep learning have revolutionized almost all stages of drug discovery and development^**12**^, such as identifying novel targets^**13**^, improving small-molecule compound design and optimization^**14**^, increasing understanding of disease mechanisms^**15,16**^, and developing new biomarkers for prognosis^**17,18**^. Therefore various deep learning methods for antiviral drug prediction have been proposed, including MMFA-DTA^**19**^, MT-DTI^**20,21**^, GDL^**22**^, and COVIDVS^**23**^. Moreover, numerous methods for drug-target interaction (DTI) prediction, such as DeepCPI^**24**^, DeepDTA^**25**^, GraphDTA^**26**^, DeepConvDTI^**27**^, TransformerCPI^**28**^, and MolTrans^**29**^, though not specifically designed for antiviral drug discovery, can also be applied to predict antiviral drugs by targeting virus-related proteins. However, existing methods have several limitations: (1) They can only predict antiviral compounds against specific virus, which are unable to identify compounds with broad-spectrum antiviral activity. (2) Evaluating a compound’s antiviral potential only based on ligand-target affinity is not enough, lacking predictions from the perspective of antiviral phenotypes when target information is unavailable. (3) These models lack interpretability, failing to identify which atom or functional group in compounds, as well as which binding site on targets, greatly influence antiviral efficacy.

Here, we proposed DeepAVC, a framework for highly accurate broad-spectrum antiviral compound prediction from the vast chemical space, including two models, DeepPAVC for phenotype-based prediction and DeepTAVC for target-based prediction. As a result, DeepAVC substantially outperforms existing models in predicting antiviral compounds and enables comprehensive prediction at both the phenotype and target levels. Building on this, we applied the DeepPAVC model for virtual screening of potential antiviral compounds across 10 large-scale compound libraries and utilized the DeepTAVC model to predict potential antiviral compounds targeting the HIV reverse transcriptase. Furthermore, we selected 30 predicted potential antiviral compounds for wet-lab experimental validation, identifying MNS and NVP-BVU972 as two novel compounds with excellent broad-spectrum antiviral activity. Next, we used the DeepTAVC model to analyze key functional groups of antiviral compounds and important binding sites on targets, demonstrating the high interpretability of our model. For user convenience, we have developed a user-friendly website (http://www.cuilab.cn/DeepAVC) designed to accelerate the discovery and development of antiviral drugs.

## Result

### Overview of DeepAVC

A typical retrovirus life cycle mainly consists of nine stages **(Fig. 1a)**. Targeting key viral proteins involved in these stages allows for the development of drugs with distinct antiviral mechanisms, named target-based antiviral compound prediction. In contrast, when target information is unavailable, it becomes necessary to predict antiviral activity only based on the compound’s structure, called phenotype-based antiviral compound prediction. Given that, DeepAVC comprises two models, DeepPAVC for phenotype-based prediction and DeepTAVC for target-based prediction. We collected the experimental measured antiviral-related bioactivity data based on cell assays from the ChEMBL^**30,31**^ database to construct the PAVC dataset for training DeepPAVC model. As for DeepTAVC, we first integrated compound-protein binding affinity data from ChEMBL and BindingDB^**32**^ to build the CPI dataset for the first stage of model training. We then refined this dataset to focus on antiviral-related data, creating the TAVC dataset, which was used for the second stage of DeepTAVC training **(Fig. 1b & Methods)**. The DeepPAVC model only takes compound information as input, utilizes a pre-trained molecular encoder to extract compound features, and outputs the antiviral activity score of the input compound **(Fig. 1c & Methods)**. In contrast, the DeepTAVC model requires input from two modalities: compounds and proteins. It employs two distinct pre-trained encoders to extract features from each modality separately, then captures intra- and inter-modality interaction patterns through self-attention and cross-attention mechanisms, and outputs the interaction score between the compound and the protein **(Fig. 1d & Methods)**. We utilized DeepAVC for virtual screening of compound libraries and validated the screening results through literature evidence, molecular docking, and wet-lab experiments. Furthermore, we discovered that the DeepPAVC and DeepTAVC models can be used synergistically: DeepPAVC can perform an initial screening of compounds for antiviral activity, and then DeepTAVC can predict the targets of antiviral drugs, providing insights into their antiviral mechanisms. Lastly, the DeepAVC framework demonstrated strong interpretability by identifying key functional groups of antiviral drugs and important binding sites on their targets **(Fig. 1e)**.

**Fig. 1:**
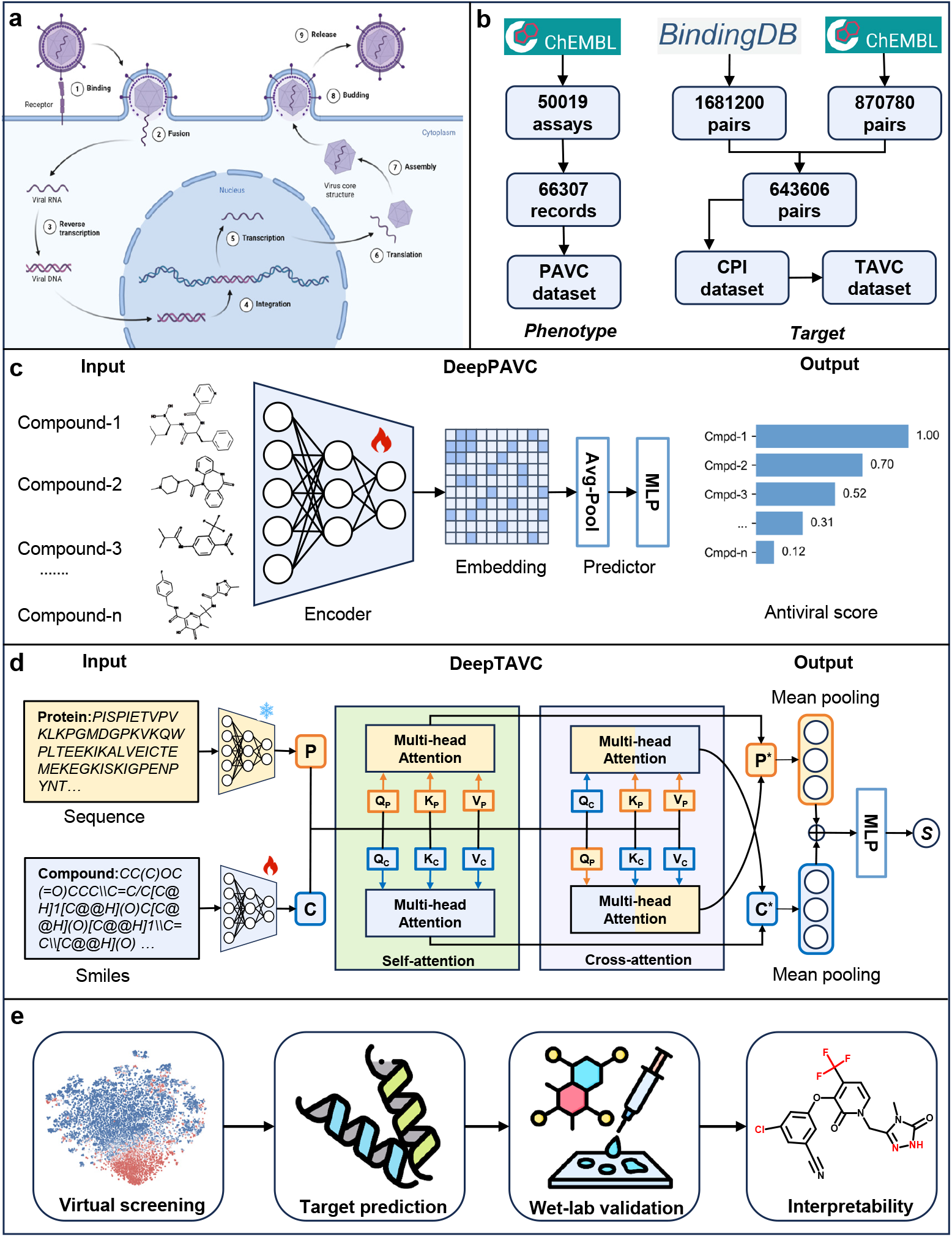
Overview of the workflow. **(a)** The typical replication process of retrovirus. **(b) Dataset construction**: The PAVC dataset for training the DeepPAVC model was built based on the compound antiviral activity data collected from the cell-based assays in the ChEMBL database. The compound-protein binding affinity data were collected from the ChEMBL and BindingDB database to construct the CPI dataset, and then selected the antiviral-related data to build the TAVC dataset for training the DeepTAVC model. **(c) Model architecture of DeepPAVC**: The DeepPAVC model only takes the compound information as input, utilizes a pre-trained molecular encoder to extract compound features, and outputs the predicted antiviral activity score of the input compound. **(d) Model architecture of DeepTAVC:** The DeepTAVC model needs both compounds and proteins information as inputs, which employs two distinct pre-trained encoders to extract features from each modality separately. It captures intra- and inter-modality interaction patterns through self-attention and cross-attention mechanisms, and outputs the interaction score between the input compound and the protein. **(e)** The main applications and analyses of the DeepAVC, including virtual screening of novel antiviral compounds, predicting the potential targets of antiviral compounds, wet-lab experimental validation of the predicted antiviral compounds and high model interpretability of identifying key functional groups of antiviral compounds.

### Performance evaluation and comparison of DeepAVC

The performance of the DeepPAVC model depends on the molecular encoder. Therefore, we first evaluated the performance of various molecular representation methods, including molecular fingerprints and descriptors generated by RDKit^**33**^, as well as distinct pre-trained encoders for 1D^**34-36**^, 2D^**37**^, and 3D^**38**^ representations of compounds (**Supplementary Table 1**). We used area under the receiver operating characteristic curve (AUROC) and area under the precision-recall curve (AUPRC) as evaluation metrics. The results showed that the KPGT model^**37**^, based on 2D molecular graph encoding of compound structures, achieved the best performance, with an AUROC of 0.91 and an AUPRC of 0.90 (**Fig. 2a & Supplementary Fig. 1**), so we selected KPGT as molecular encoder for the DeepPAVC model.

**Fig. 2:**
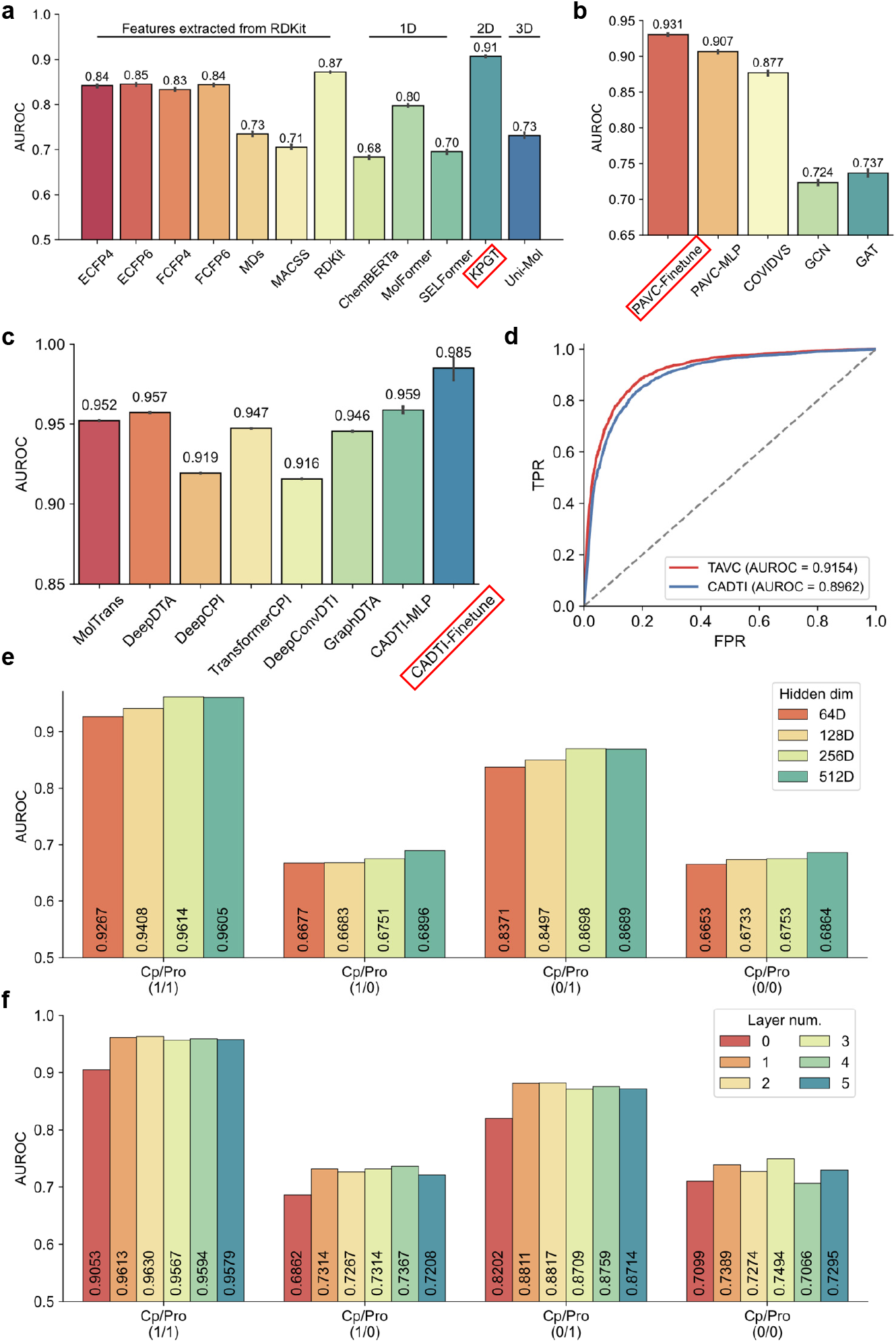
Performance presentation and comparison of DeepAVC models. **(a)** Performance comparison of different molecular representations in antiviral drug prediction tasks. **(b)** Performance comparison between the DeepPAVC model and other baseline models. **(c)** Performance comparison between the DeepTAVC model and other baseline models. **(d)** Performance comparison between the model (DeepTAVC) trained with antiviral-related datasets in the second training stage and the model trained without them (CADTI). **(e)** Performance comparison of DeepTAVC models with different hidden layer dimensions across four distinct test sets. **(f)** Performance comparison of DeepTAVC models with varying attention layer numbers across four different test sets. AUROC: Area under the ROC curve. 1D: one-dimensional molecular representation. 2D: two-dimensional molecular representation. 3D: three-dimensional molecular representation.

Subsequently, we compared the performance of the DeepPAVC model with other baseline models. We identified COVIDVS^**23**^ as a relevant comparator because it is currently the only open-source model that predicts antiviral activity based solely on the compound information. Additionally, we trained two baseline models, based on GCN and GAT, which utilize 2D molecular graph encodings of compound structures as well. Moreover, it has been reported that fine-tuning all trainable parameters of pre-trained large models outperforms the combination of pre-trained models with frozen parameters equipped with classification heads^**39,40**^, so we also compared the performance of the above two training strategies of the KPGT model. The results showed that the DeepPAVC model outperformed all baseline models, achieving an AUROC value over 0.9. Furthermore, fine-tuning of the DeepPAVC model demonstrated better performance compared to the non-fine-tuning approach, with AUROC values of 0.931 and 0.907, respectively (**Fig. 2b)**.

For the DeepTAVC model, existing DTI prediction models are not specifically tailored to antiviral targets but rather encompass targets associated with various disease types. To address this, we selected a widely-used external dataset^**41,42**^ for model training and performance comparison, referring to our model trained on this dataset as CADTI (**Methods**). The results demonstrated that the fine-tuning version of CADTI achieved the best performance, with an AUROC of 0.985 and an AUPRC of 0.981, followed by the CADTI**+**MLP combination, which attained an AUROC of 0.959 and an AUPRC of 0.947. These findings highlight the advantages of our proposed model architecture in the DTI prediction task (**Fig. 2c & Supplementary Fig. 2)**.

Next, we retrained the CADTI model using our constructed CPI dataset (**Methods**), which includes more diverse compound-target affinity data. This dataset enabled the model to learn general patterns of interactions between compounds and proteins, thereby improving its generalization performance. We then conducted a second-stage training of the CADTI model using the TAVC dataset (**Methods**), resulting in the final DeepTAVC model. The results showed that the DeepTAVC model slightly outperformed the CADTI model, with AUROC values of 0.9154 and 0.8962, respectively. This improvement underscores the importance of domain-specific data for fine-tuning the model in the second training stage (**Fig. 2d)**.

We next investigated whether increasing the number of parameters in the DeepTAVC model could further enhance its performance. To comprehensively evaluate the model, we divided the test set into four scenarios based on whether the compounds and proteins were present in the training set (**Methods**). We first explored the impact of the hidden layer dimension in the attention layers. The results showed that increasing the hidden layer dimension had minimal effect on performance across all four test scenarios (**Fig. 2e**). Next, we examined the effect of the number of attention layers. The results indicate that adding an attention layer greatly enhances model performance, demonstrating that the cross-attention mechanism we employed effectively captures interaction patterns between compounds and proteins. However, further increasing the number of attention layers had little impact on model performance across four test scenarios (**Fig. 2f**). Given that, we set the dimension of the attention layer to 256 and the number of attention layer to 1.

### Virtual screening of antiviral compounds by using DeepPAVC

One of the main applications of DeepAVC is the rapid virtual screening of antiviral compounds from the vast chemical space. The DeepPAVC model, requiring only compound information as input, enables rapid high-throughput virtual screening for potential antiviral compounds. We selected 10 representative compound libraries for virtual screening (**Supplementary Table 2**), with the number of compounds ranging from 10^4^ to 10^9^ (**Fig. 3a**). These libraries encompass diverse types of compounds. For example, the DrugBank database^**43**^ mainly includes FDA-approved drugs, while the MCE database consists of commercial compounds with biological activities. The virtual screening results from the DrugBank database were visualized using the t-distributed Stochastic Neighbor Embedding (t-SNE) method for dimensionality reduction^**44**^. The results revealed that antiviral drugs and non-antiviral drugs clustered into two distinct groups, indicating that the DeepPAVC model effectively distinguishes between antiviral and non-antiviral compounds. The top four compounds ranked by prediction scores were Beclabuvir, Cabotegravir, Ziresovir, and Raltegravir, all of which are either FDA-approved or in clinical trial stages as antiviral drugs (**Fig. 3b**). Similarly, the screening results from the MCE database exhibited comparable patterns. The top four compounds identified were ST-193, Dolutegravir, Roblitinib, and HIV-1 inhibitor-8 (**Fig. 3c**), demonstrating that DeepPAVC not only identifies known antiviral drugs but also highlights novel candidate compounds with potential antiviral activity. The virtual screening results for the remaining eight databases were also provided (**Extended Data Fig. 1 & Supplementary Table 3)**.

**Fig. 3:**
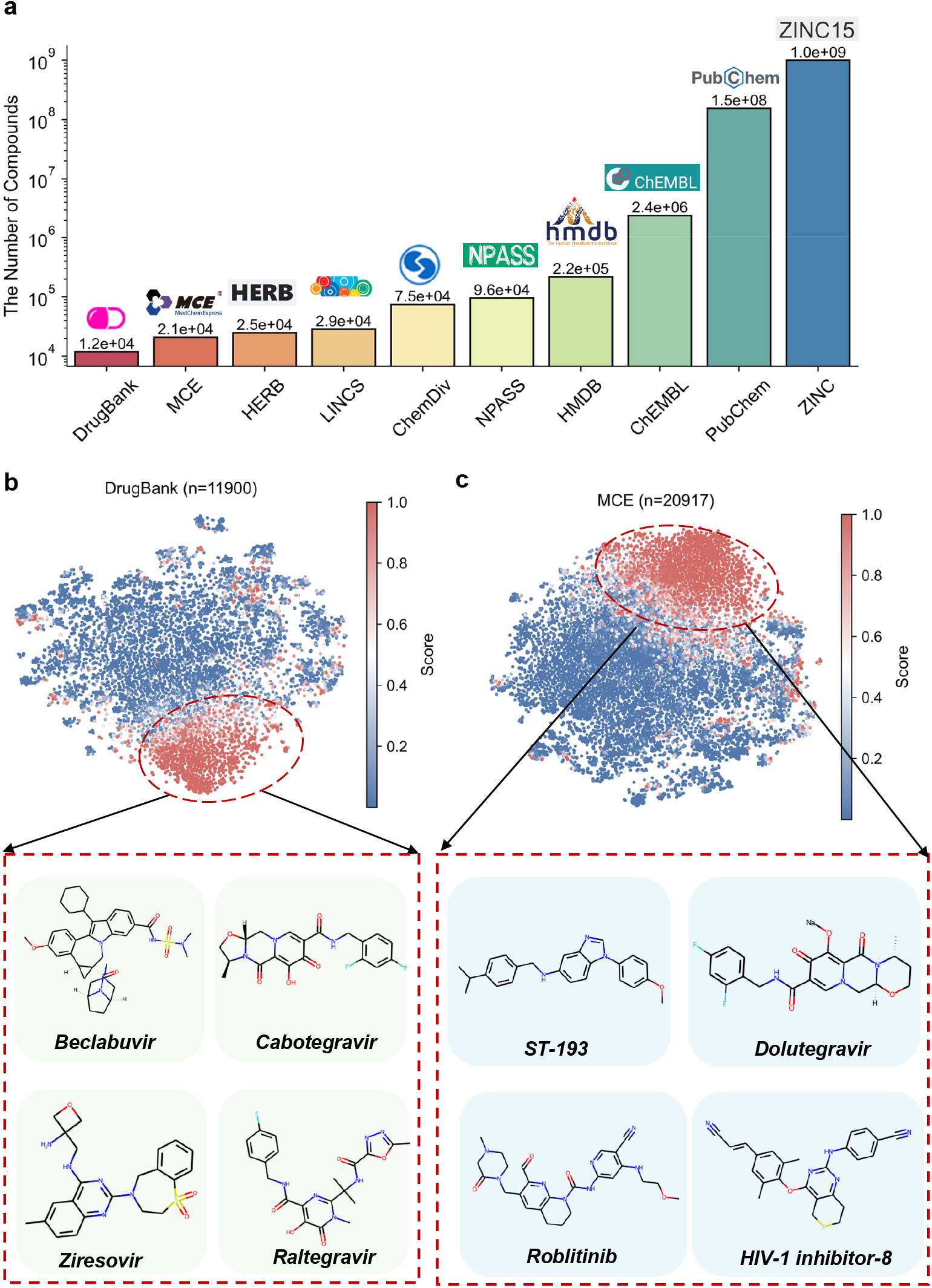
Virtual screening of broad-spectrum antiviral compounds using the DeepPAVC model. **(a)** The number of compounds in 10 large-scale chemical databases used for virtual screening. **(b)** Virtual screening results from the DrugBank database and structures of the top 4 compounds ranked by predicted antiviral scores, including Beclabuvir, Cabotegravir, Ziresovir, and Raltegravir. **(c)** Virtual screening results from the MCE database and structures of the top 4 compounds ranked by predicted antiviral scores, including ST-193, Dolutegravir, Roblitinib, and HIV-1 inhibitor-8.

### Virtual screening of antiviral compounds by using DeepTAVC

DeepTAVC enables target-based virtual screening of antiviral compounds when interested target information is available. HIV is a highly lethal virus that poses a severe threat to human health^**45**^. As a typical retrovirus, HIV undergoes a life cycle involving nine stages (**Fig. 1a**), with reverse transcriptase being one of the key therapeutic targets for treating HIV infection. Therefore, we selected the HIV-1 reverse transcriptase (Uniprot ID: Q72547) as the interested target and performed target-based virtual screening for anti-HIV compounds. Using DeepTAVC, we first screened compounds from the DrugBank database, identifying drugs with the highest prediction scores, including Etravirine, Doravirine, Capravirine, Rilpivirine, and Dapivirine (**Fig. 4a**). These compounds are either FDA-approved or in clinical trial stages as non-nucleoside reverse transcriptase inhibitors, demonstrating that DeepTAVC successfully identifies known inhibitors with anti-HIV reverse transcriptase activity. Next, we extended the virtual screening using the ChEMBL database to discover unreported compounds targeting HIV-1 reverse transcriptase. To validate the model’s screening effectiveness, we retrieved the structure of the HIV-1 reverse transcriptase complexed with a known small-molecule inhibitor from the PDB database (PDB ID: 2JLE^**46**^)^**47,48**^. We then selected the top five anti-HIV compounds predicted by DeepTAVC, and compared the binding energies of these compounds with the original ligand in the HIV-1 reverse transcriptase complex using molecular docking (**Fig. 4b & Methods**). The results showed that the binding energies of the top five compounds predicted by DeepTAVC, ranging from -11.3 kcal/mol to -12.5 kcal/mol, substantially better than the original ligand’s binding energy of -9.6 kcal/mol, indicating that DeepTAVC can identify HIV-1 reverse transcriptase inhibitors with higher binding affinity (**Fig. 4c**). Molecular docking interaction diagrams further revealed that these high-affinity compounds screened by DeepTAVC exhibited enhanced interactions with amino acid residues within the active site of HIV-1 reverse transcriptase. For example, CHEMBL4475543 formed additional hydrogen bonds with HIS-235 and TYR-318, as well as a π-π stacking interaction with PHE-227, thereby substantially improving its binding affinity (**Fig. 4b & Fig. 4c**).

**Fig. 4:**
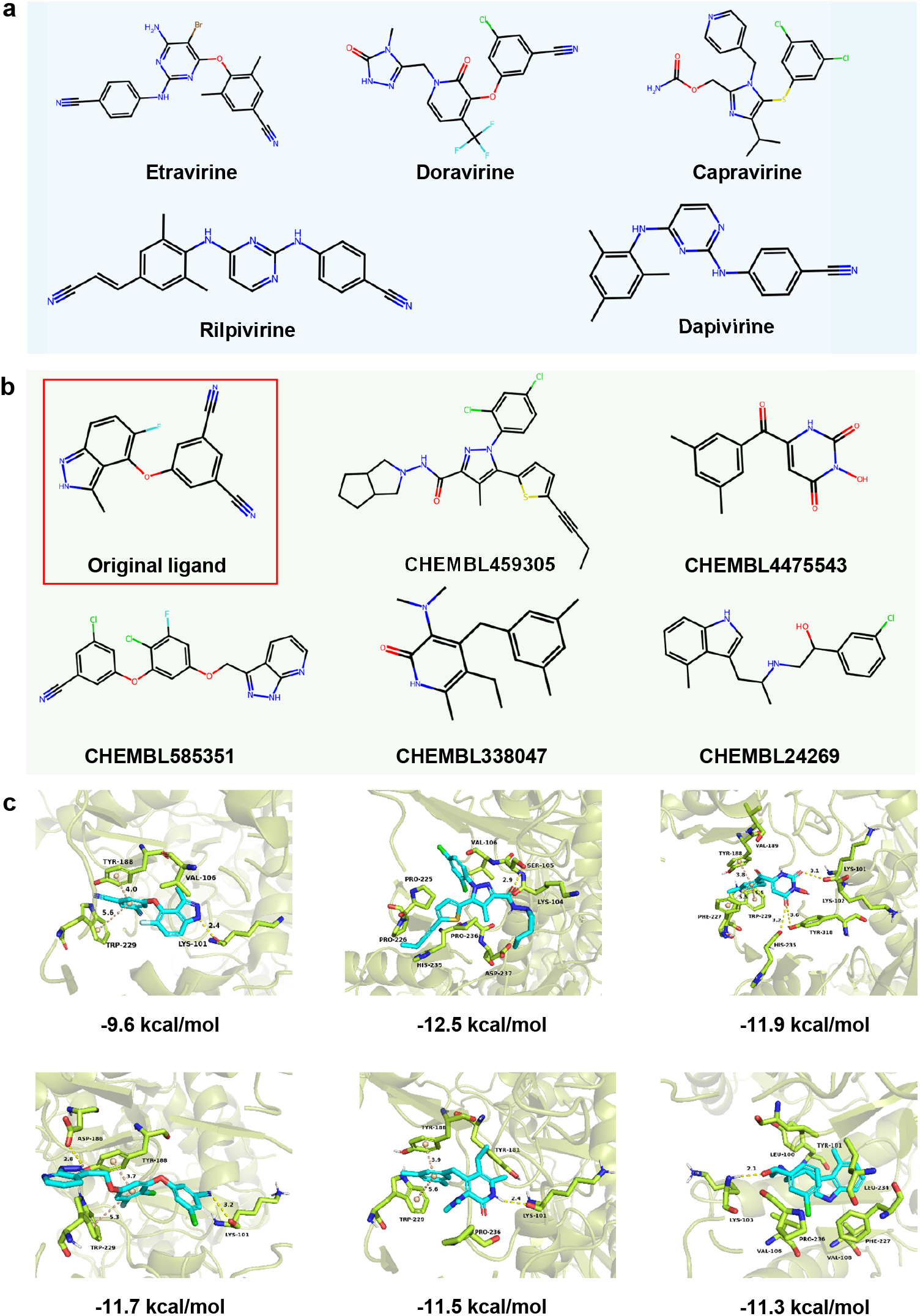
Virtual screening of anti-HIV compounds using the DeepTAVC model. **(a)** Top 5 predicted compounds from the DrugBank database with potential high binding affinity to HIV-1 reverse transcriptase. **(b)** The original antiviral compound (CHEMBL473119) of the HIV-1 reverse transcriptase (PDB ID: 2JLE) and the top 5 predicted compounds from the ChEMBL database with potential higher binding affinity. **(c)** Comparison of binding affinities between compounds screened by DeepTAVC from the ChEMBL database and the original antiviral compound (CHEMBL473119) of the HIV-1 reverse transcriptase using molecular docking.

### Experimental validation of the antiviral activity of compounds identified by DeepAVC

To further validate the virtual screening results, we selected 30 potential antiviral compounds screened by DeepPAVC from the MCE database for wet-lab experimental validation (**Supplementary Table 4**). Due to the outstanding performance of DeepPAVC, the top compounds with the highest predicted score are mostly known antiviral drugs or antiviral compounds supported by literature evidence. Therefore, we mainly excluded compounds with known antiviral activity and retained previously unreported compounds to find novel antiviral agents.

The antiviral efficacy of these compounds was first assessed against five distinct viruses, including DNA virus Herpes Simplex Virus 1, F strain (HSV-1 F strain), positive-strand RNA viruses Encephalomyocarditis Virus (EMCV) and Mouse Hepatitis Virus, A59 strain (MHV-A59), and negative-strand RNA viruses Influenza A Virus, PR8 strain (IAV-PR8) and Vesicular Stomatitis Virus (VSV), by measuring viral loads post-treatment via the RT-qPCR (**Methods**). As shown in **Figure 5a**, 28 out of the 30 compounds inhibited at least one virus, with six compounds demonstrating broad-spectrum antiviral activity by effectively suppressing all five tested viruses (**Fig. 5a & Supplementary Table 5**). Notably, Compound 13 (NVP-BVU972) and Compound 18 (MNS) exhibited the strongest antiviral activity among these top candidates. To further confirm the antiviral activity of Compounds 13 and 18, we selected Compound 9 (Sisunatovir) to serve as a positive control drug and assessed viral levels using VSV-GFP as a reporter virus. Microscopy analysis showed that all three compounds exhibited the antiviral activity, resulting in a substantial reduction of viral replication compared with the control group. More strikingly, Compound 18 completely eliminated viral infection in HeLa cells compared to the other two compounds (**Extended Data Fig. 2**). Furthermore, flow cytometry and western blot analyses produced similar results, providing strong evidence of the potent antiviral activity of Compound 18 in particular (**Fig. 5b-5c**).

**Fig. 5:**
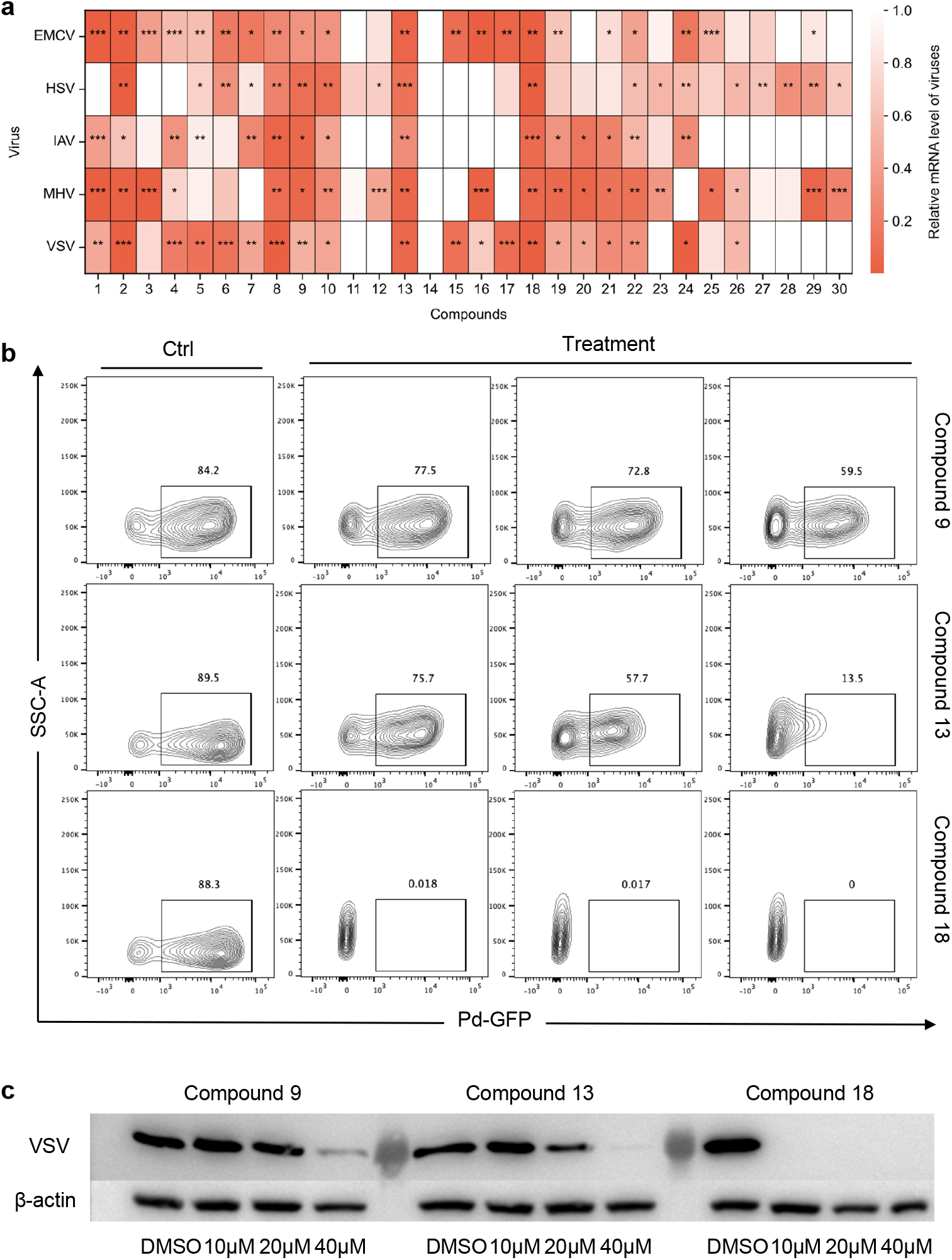
Experimental validation of the antiviral efficacy of compounds identified through DeepAVC. **(a)** Antiviral activity of 30 compounds against different virus strains. Heatmap illustrating the antiviral effects of 30 compounds against five virus strains: EMCV, HSV, IAV, MHV, and VSV. The color intensity in each cell reflects the antiviral efficacy, with darker shades indicating stronger antiviral activity. **(b)** Flow cytometry assessment of viral replication in HeLa cells treated with antiviral compounds. HeLa cells were infected with VSV-GFP for 12 hours and treated with Compound 9, Compound 13, Compound 18, or DMSO (Ctrl). Flow cytometry was used to quantify GFP fluorescence (Pd-GFP) as an indicator of viral replication. The contour plots display the distribution of GFP fluorescence intensity (X-axis) versus side scatter (SSC-A, Y-axis), with the percentage of GFP-positive cells in each quadrant, reflecting the effectiveness of each treatment in reducing viral replication. **(c)** Western blot analysis of viral replication following treatment with antiviral compounds. HeLa cells were infected with VSV-GFP for 12 hours and treated with Compound 9, Compound 13, or Compound 18 at concentrations of 10 μM, 20 μM, and 40 μM, or DMSO as a control. Western blot analysis was performed to detect GFP expression, which serves as a marker for viral protein production. β-actin was used as a loading control to ensure equal protein loading across samples.

These findings demonstrate that the compounds identified by DeepPAVC exhibit promising antiviral activity, with compound 18 even outperforming compound 9 (Sisunatovir), an antiviral drug currently in clinical trials. This highlights DeepPAVC as a powerful tool for antiviral drug discovery, with great potential for identifying novel broad-spectrum antiviral agents.

### Enhancing model interpretability in antiviral drug discovery by DeepTAVC

Although DeepPAVC enables rapid virtual screening of antiviral compounds without requiring protein information, it lacks interpretability, as the targets of the identified compounds remain unknown. DeepTAVC mitigates this limitation by predicting potential targets for the compounds identified through DeepPAVC, thereby providing insights into their antiviral mechanisms. The TAVC dataset we constructed includes 238 known antiviral-related targets (**Methods**), which cover various aspects of antiviral mechanisms, such as virus replication, virus entry, virus assembly, virus budding, host immune response, and host inflammation **(Supplementary Table 6)**. Therefore, we applied the DeepTAVC model to the 30 compounds screened by DeepPAVC from the MCE database, and predicted their binding scores for the 238 antiviral-related targets. The results revealed diverse antiviral modes of action among these compounds. For instance, Compound 2, Compound 9, and Compound 14 exhibited high affinities for almost all antiviral-related targets, suggesting they may act as broad-spectrum antiviral drugs. In contrast, Compound 13 and Compound 18 showed low affinities for all known antiviral targets, despite their strong antiviral phenotypes (**Fig. 5a**). This indicates that these two compounds might represent novel broad-spectrum antiviral drugs with entirely new mechanisms of action. For the remaining compounds, some displayed strong affinities for specific antiviral targets, suggesting they may be effective against particular types of viruses (**Fig. 6a**). These findings demonstrate that DeepPAVC can identify not only broad-spectrum and target-specific antiviral drugs but also compounds with novel antiviral mechanisms.

**Fig. 6:**
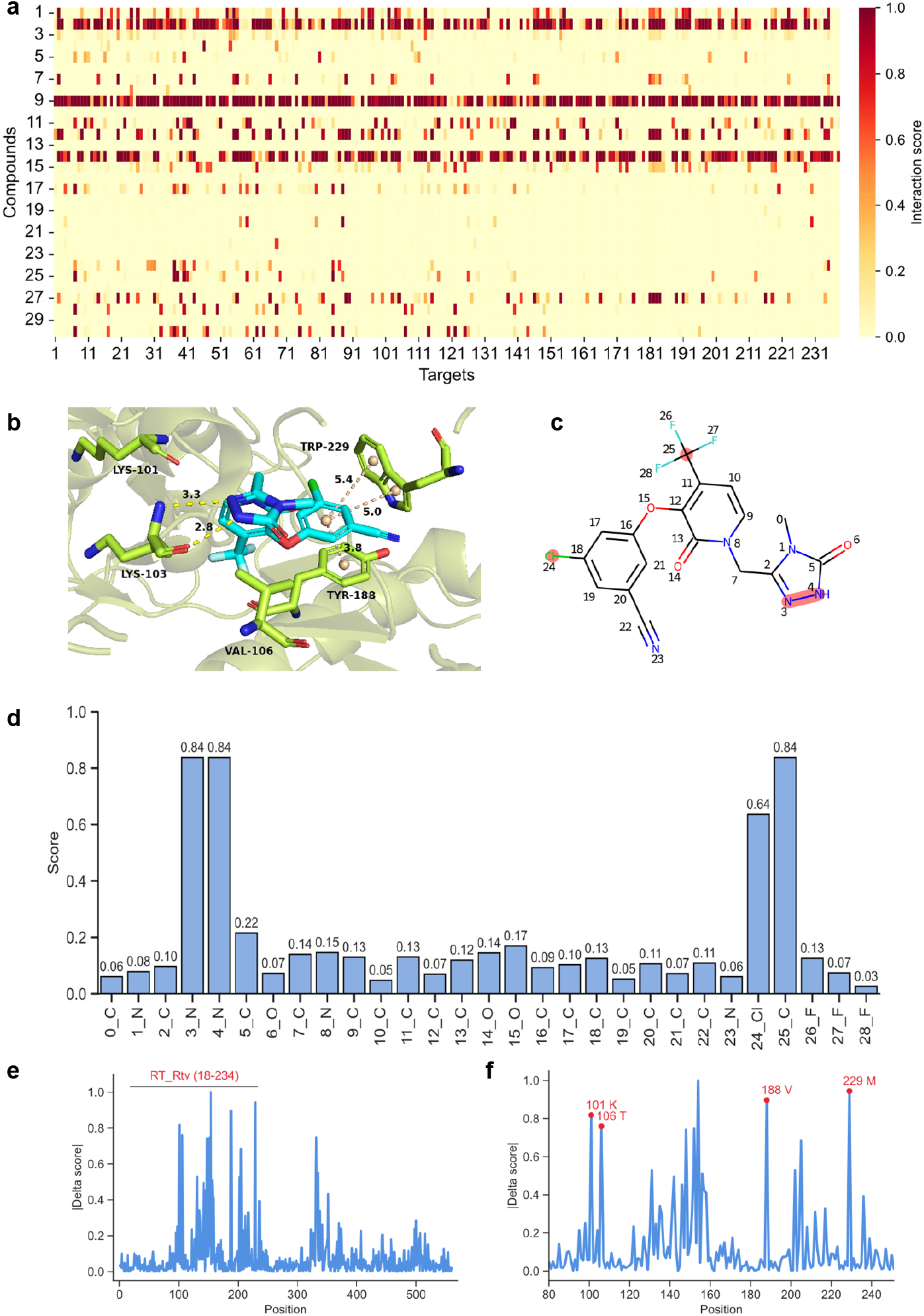
Enhancing interpretability in antiviral drug discovery by DeepTAVC model. **(a)** Predicted potential antiviral targets of 30 compounds screened by the DeepPAVC model from the MCE database using the DeepTAVC model. **(b)** Binding mode of Doravirine with HIV-1 reverse transcriptase (PDB ID: 4NCG). **(c)** Structure of Doravirine with key atoms and functional groups highlighted in red. **(d)** Atom-wise attention scores of Doravirine extracted from the DeepTAVC model, reflecting each atom’s contribution to compound-protein binding. **(e)** Relative activity change scores for each position in the HIV reverse transcriptase sequence, reflecting each site contribution to compound-protein binding. **(f)** Magnified view of the 80-250 aa region in Fig. 6e, with key amino acid positions highlighted in red. RT_Rtv (18-234): the HIV reverse transcriptase domain, a key region for its enzymatic activity, is located at positions 18-234 in the sequence. 101K: Lysine at position 101. 106T: Threonine at position 106. 188V: Valine at position 188. 229M: Methionine at position 229.

We further utilized the DeepTAVC model to explore the specific binding patterns of antiviral drugs with their targets. As a case study, we selected the complex structure of Doravirine, an orally bioavailable non-nucleoside reverse transcriptase inhibitor, bound to HIV reverse transcriptase (PDB ID: 4NCG^**49**^). Doravirine forms multiple interactions with amino acid residues in the active pocket of the target, enhancing its binding affinity (**Fig. 6b & Fig. 6c**). Using the DeepTAVC model, we extracted attention maps representing compound-protein interactions and normalized the attention scores between 0-1 for Doravirine. These attention scores highlight the importance of specific atoms or functional groups in Doravirine during its interaction with HIV reverse transcriptase, revealing that the 3rd and 4th nitrogen (N) atoms, the 24th chlorine (Cl) atom, and the 25th carbon (C) atom in Doravirine had significantly higher attention scores compared to other positions (**Fig. 6c-d & Supplementary Fig. 3**). In fact, the 3rd and 4th nitrogen atoms form hydrogen bonds with LYS-103 of HIV reverse transcriptase, while the 24th chlorine atom and 25th carbon atom correspond to a trifluoromethyl functional group, which notably enhances the hydrophobicity of the compound. This increased hydrophobicity allows Doravirine to better access the hydrophobic active pocket of the target, thereby improving its binding affinity. These findings demonstrate that the attention maps generated by DeepTAVC effectively identify the key atoms and functional groups involved in drug-target interactions.

Finally, we investigated whether DeepTAVC could also provide interpretability from the perspective of the target. We computed the relative activity change score for each position in HIV reverse transcriptase (**Methods**). This score measures the importance of each amino acid residue in compound-protein binding, with higher absolute values indicating greater importance. We observed that the RT_Rtv domain ranging from 18 to 234 within the protein sequence, which is critical for HIV reverse transcriptase activity, exhibited significantly higher mutation activity scores compared to other regions (**Fig. 6e**). Furthermore, among the amino acids near the Doravirine binding pocket, mutations at K-101, T-106, V-188, and M-229 caused substantial changes in predicted binding activity score, identifying these residues as key binding sites for Doravirine (**Fig. 6f**). Actually, the phenyl ring of Doravirine forms π-π stacking interactions with TYR-188 and TRP-229 of HIV reverse transcriptase, highlighting that the relative activity change score calculated by DeepTAVC accurately reflect the key binding residues in protein targets during compound-protein interactions. These results indicate that DeepTAVC can substantially enhance model interpretability in antiviral drug discovery.

## Discussion

Viral infections present a major challenge to human health, underscoring the urgent need for the development of effective antiviral drugs to protect lives. Here, we propose DeepAVC, a deep learning framework based on pre-trained large models, comprising DeepPAVC and DeepTAVC, which predict potential antiviral compounds at the phenotype-level and target-level, respectively. The DeepAVC framework addresses two scenarios in antiviral drug discovery: (1) When target information is unavailable or irrelevant, the DeepPAVC model predicts antiviral compounds only based on compound information. (2) When target information is available, the DeepTAVC model utilizes both compound and protein information for prediction. By integrating these approaches, DeepAVC offers a versatile and comprehensive solution for antiviral drug discovery under diverse research conditions. Moreover, the DeepPAVC and DeepTAVC models are complementary to each other. When potential antiviral compounds are initially identified using the DeepPAVC model, the DeepTAVC model can subsequently predict their potential target interactions, providing insights into their antiviral mechanisms. Conversely, when compounds capable of binding to a target of interest are first identified using the DeepTAVC model, the DeepPAVC model can be employed to further assess their antiviral activity. This additional step is crucial for eliminating false-positive compounds that may bind to the target but lack the corresponding biological activity. By integrating these two models, researchers can achieve a more accurate and comprehensive antiviral compound discovery process.

Although DeepAVC enables rapid and accurate screening of antiviral compounds from the vast chemical space, it has certain limitations. First, the training data for DeepPAVC is solely derived from ChEMBL, which may introduce biases and limit the model’s generalizability. Second, the DeepTAVC model considers only the sequence information of target proteins, excluding structural information, which could impact prediction accuracy. Lastly, the optimal integration of DeepPAVC and DeepTAVC for more in-depth and effective antiviral drug discovery remains an area for further exploration. Future explorations and improvements in this domain include the following: (1) Incorporating structural information of proteins: Enhancing the predictive performance of the model by integrating structural data when predicting drug-target interactions. (2) Expanding training datasets: Collecting a broader range of antiviral phenotypic experimental data to reduce dataset biases and improve the model’s generalizability. (3) Accounting for confounding factors: Considering the effects of factors such as compound dosage and treatment duration on the performance of antiviral drugs.

## Methods

### Dataset construction

#### The PAVC dataset

In this study, we select the ChEMBL_34 database to construct PAVC dataset for training DeepPAVC model. The ChEMBL_34 database was released in March 2024 and contains 2,496,335 distinct compounds, 16,003 targets, 1644390 assays, 21,123,501 activities, and 92,121 source publications.

#### Data filtering

We first downloaded all assays available in ChEMBL and filtered out virus-related assays using three strategies: **(1) Organism-based filtering**: we selected assays with the organism term categorized as “virus”, resulting in 43,843 virus-related assays. **(2) Keyword-based filtering**: we extracted assays containing the keyword “virus” in the Description field, identifying 42,140 virus-related assays. **(3) Virus-specific filtering**: we selected 44 common viruses and extracted assays that mentioned their names in the Description filed, obtaining 36,189 assays. Next, we combined the assay sets obtained from these three approaches by tasking the union based on their unique assay IDs, resulting in 64,524 assays. After excluding assays unrelated to antiviral activity measurements, we finalized a dataset comprising 50,019 virus-related assays.

#### Data processing

We next processed the virus-related bioactivity dataset using the following step: (1) The assay type was set to ‘F’, which represents the data measuring the biological effect of a compound to cell. (2) Activity data with metrics of inhibition constant (*k*_*i*_), dissociation constant (*k*_*d*_), half-maximal inhibitory concentration (*IC*_50_), and half-maximal effective concentration (*EC*_50_) are reserved, while activity data without bioactivity records or whose label could not be unequivocally assigned (e.g., activity < 100 *μM* or activity > 1 *μM*) are removed. (3) The units of bioactivity data (i.e. *g* / *ml,M, μM*) are converted into the standard unit in *nM*. (4) If a compound has multiple biological activity values, then we calculated the average of them as the final value. (5) We transformed *k*_*i*_, *k*_*d*_, *IC*_50_, and *EC*_50_ to *pK*_*i*_, *pK*_*d*_, *pIC*_50_, and *pEC*_50_, and then selected the threshold of 6 to define positive (with antiviral activity) and negative data (without antiviral activity) because only compounds whose *IC*_50_, *EC*_50_, *k*_*i*_, and *k*_*d*_are at the *μM* or even *nM* level will be further optimized^**50**^. (7) Compounds with only positive or negative labels were removed. (8) We obtained the SMLIES representations of all compounds using RDKit, and those with the number of atoms higher than 90 were excluded.

Finally, we constructed the PAVC dataset including 55861 distinct compounds, 26626 positive samples and 29235 negative samples in total.

#### Compound-Protein Interaction (CPI) dataset

To collect comprehensive data of compound-protein interactions, we integrated compound-protein binding affinity data from the ChEMBL database and the BindingDB database. The BindingDB database contains approximately 3.0 million data points for 1.3 million compounds and 9,500 targets.

#### The ChEMBL database

We filtered and processed the measured binding affinity data from the ChEMBL database using the following procedure: (1) The target type was set to ‘SINGLE PROTEIN’, and the molecule type was set to ‘Small molecule’; (2) Data with a confidence score of 9 were retained; (3) The assays with assay type of ‘B’, representing data measuring binding of the compound to a molecular target, were reserved; (4) Data with metrics of *k*_*i*_, *k*_*d*_, *IC*_50_, and *EC*_50_ were selected, and the units of these data were converted into the standard unit in *nM*; (5) For a molecule with multiple bioactivity values, the final binding affinity value was generated by averaging the available bioactivity records; (6) We transformed *k*_*i*_, *k*_*d*_, *IC*_50_, and *EC*_50_ to *pk*_*i*_, *pk*_*d*_, *pIC*_50_, and *pEC*_50_, respectively; (7) The threshold of 6.5 was selected to define positive (with binding activity) and negative data (without binding activity). (8) Data whose atom number was higher than 90 or whose protein sequence length exceeded 4000 were filtered out.

#### The BindingDB database

We employed the same data processing pipeline as the ChEMBL database, as described above.

Finally, we kept the unique pairwise compound-target samples from the processed ChEMBL database and the BindingDB database, resulting in the CPI database including 198,246 distinct compounds, 6193 distinct proteins, 321534 positive samples and 321945 negative samples.

#### The TAVC dataset

Based on the CPI dataset, we further constructed the TAVC dataset to train the DeepTAVC model, enabling target-based antiviral compound prediction. To be specific, we first collected approximately 200 antiviral-related target proteins from the therapeutic target database (TTD)^**51**^ and other literature sources. These proteins were well-annotated with their names, uniprot ids, sequences, functions, and antiviral mechanisms (**Supplementary Table 6**). Next, we filtered the CPI dataset, retaining only samples associated with these antiviral targets, resulting in the TAVC dataset, which contains 55,452 positive samples and 61,426 negative samples.

#### The external CPI dataset

To compare the performance of our DeepTAVC model with existing DTI prediction baseline models, we selected a widely-used benchmark dataset^**41,42**^, which includes 14,636 compounds, 2,623 proteins, 20,674 positive samples, and 28,525 negative samples.

#### Compound representation

In this study, we utilize a SMILES representation for a small molecule and abstract it as molecular graph *G* = (*V,E*), where 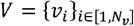 stands for the set of nodes (i.e. atoms of a molecule), and *N*_*v*_ represents the number of nodes. The features of nodes and edges in the molecular graph are initialized via RDKit. We employ the same intial features of nodes and edges as those used in KPGT. We then use KPGT to extract the learned embedding of each node, denoted as 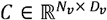, where *D*_*v*_ stands for the dimensions of embeddings of nodes.

#### Protein representation

Here, we represent a protein sequence as 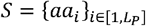, where *S* denotes a protein sequence, *aa*_*i*_ represents the *i-*th amino acid in *S*, and *L*_*s*_ is the length of *S*. We next utilize ESM-2^**52**^, a widely-used protein language model, to encode each *aa* within *S* into a high-dimension embedding, denoted as 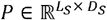, where *D*_*s*_ stands for the dimensions of embeddings of amino acids.

#### Multi-head attention (MHA)

After obtaining the compound feature *C* and the protein feature *P*, we utilize a multi-head attention mechanism to capture the complex interactions between them:

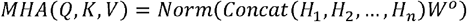

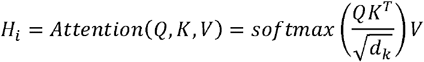

Where *MHA* denotes multi-head attention, *H*_*i*_ stands for the *i-*th attention head, *Q, K*, and *V* represent query, key, and value matrix respectively, 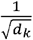 is the scaling factor of the dot-product attention, *T* denotes the matrix transposition, *Norm* refers to the layer normalization operation. *Concat* denotes the concatenation operation. *W*^0^ represents the weight matrix for output transformation.

#### Model architecture

**DeepPAVC**. DeepPAVC requires only the SMILES representations of compounds as input. Here, we utilize KPGT as a molecule encoder 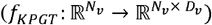 to extract the atom-level compound feature 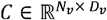, and then employ a pooling aggregation strategy to obtain the overall molecular representation. Finally, we apply a MLP to output the antiviral score of input compounds. This process is delineated by the following equation:

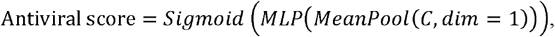

where *MeanPool* calculates the element-wise mean of all tokens across the sequence dimension, and *MLP* denotes a multi-layer perceptron, 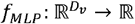 that maps the embedding to a scalar value. **DeepTAVC**. DeepTAVC requires both the SMILES of the compound and the amino acid sequence of the protein as inputs. Here, we first employ KPGT 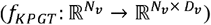 and ESM-2 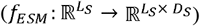 to extract the compound features *C* and protein features *P*, respectively. Next, we employ the MHA mechanism to refine *C* and *P*, including a self-attention mechanism to capture the interaction patterns within compounds (atoms) and within protein (residues), as well as a cross-attention mechanism to model the cross-modal interactions between compounds (atoms) and proteins (residues).

To be specific, we first employ two distinct linear projection layers to map the compound features *C* and protein features *P* into the same dimensional space:

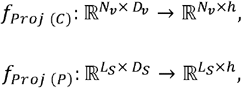

where h denotes the unified dimensions of embeddings used for MHA mechanism.

Next, we use two distinct sets of projection matrices for compounds and proteins to construct the query, key and value matrices for the MHA mechanism:

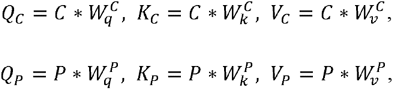

where 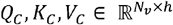 and 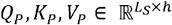 are the queries, keys and values for compound and protein, respectively, the projection matrices 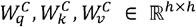 and 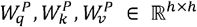 are used to derive the queries, keys and values, respectively.

Then, we apply the multi-head, self-attention and cross-attention mechanisms to refine the representations of each atom (compound) and residue (protein) as below:

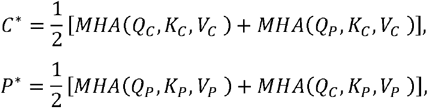

where 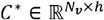 and 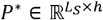 are the updated embeddings of the compound and protein obtained through the MHA mechanism.

After that, we utilize a pooling aggregation strategy for both *C*^*^ and *P*^*^ independently and then concatenate them to obtain the final joint representation *F*:

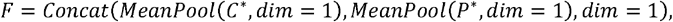

where *MeanPool* calculates the element-wise mean of all tokens across the sequence dimension, and *Concat* denotes the concatenation operation of the resulting mean vectors.

Finally, we apply a MLP to output the affinity score between the input compound and protein. This process is delineated by the following equation:

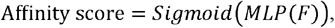

where *F* denotes the joint representation for the input compound and protein obtained through the MHA mechanism.

### Training strategy

#### DeepPAVC

We split the PAVC dataset into training, validation, and test sets in an 8:1:1 ratio. The model parameters of the molecular encoder, KPGT, were set to be fully trainable. An MLP was used to output the likelihood of the input being an antiviral compound, and the model was trained by minimizing the cross-entropy loss between the predicted probabilities and the true labels. The hyperparameters of the DeepPAVC were determined through hyperparameter optimization.

#### DeepTAVC

For the DeepTAVC model, we applied a two-stage training strategy. In the first stage, the model was trained on the CPI dataset to learn broader compound-protein interaction patterns. Subsequently, in the second stage, the model was fine-tuned using the TAVC dataset to achieve optimal performance in the antiviral context. During training, all parameters of the compound encoder (KPGT) were set to be trainable, while the parameters of protein encoder (ESM-2) were frozen to ensure that the model’s parameter size could fit within GPU memory constraints. The training objective of the DeepTAVC model was to minimize the cross-entropy loss between the predicted interaction probabilities and the true labels. The hyperparameters of the DeepTAVC were determined through hyperparameter optimization.

#### Training details

All models in this study were implemented in PyTorch^**53**^ version 1.10.0 with CUDA version 11.3 and Python 3.7. An Adam optimizer with weight decay 1e-6 and learning rate 3e-5 was used to optimize the model. The model was trained with a batch size of 32 for a total of 10 epochs. Both DeepPAVC and DeepTAVC had around 100 million trainable parameters. Training was performed on one Nvidia A100 GPU.

#### Model performance comparison

All the baseline models for DeepPAVC comparison were trained on the PAVC dataset, while the performance comparison between DeepTAVC and their baseline models was conducted on the external CPI dataset. For DeepPAVC, the dataset was split into training and testing sets at a 9:1 ratio, followed by 10-fold cross-validation. In contrast, for DeepTAVC, we categorized the test sets based on the presence of compounds and proteins in the training set, resulting in four distinct subsets: (1) both compounds and proteins are seen, (2) compounds are seen but proteins are unseen, (3) compounds are unseen but proteins are seen, and (4) both compounds and proteins are unseen. This categorization enables a more comprehensive evaluation of DeepTAVC’s performance.

#### Molecular docking

In this study, the crystal structure of the HIV-1 reverse transcriptase was downloaded from the RCSB Protein Data Bank database (PDB ID: 2JLE) in PDB format, and modified using the Autodock tools 1.5.6 software. The target protein was processed through water removal, hydrogen addition, amino acid optimization and patching. The compounds used for docking were downloaded from the PubChem database^**54**^ in SDF format. The compounds were imported into Autodock tools, and all flexible keys were rotatable by default and then saved in pdbqt format, as docking ligand. Finally, Autodock Vina 1.2 was used for docking, and PyMol 3.1 was used to visualize the docking results.

#### Mutation activity change score calculation

We downloaded the ligand-receptor complex from the PDB database (PDB: 4NCG) for this case study, which is the structure of doravirine (ligand) and HIV reverse transcriptase (receptor). We first input the wild-type protein and compound into DeepTAVC to calculate the original prediction interaction score, denoted as *s*. Next, we mutated each amino acid of the protein sequence to all 20 amino acids (including itself) and calculated the prediction score *s*′. Finally, we defined the activity change score Δ*S* ∈ ℝ ^*l*× 20^ as below:

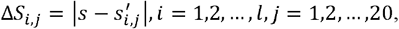

where *i* refers to the *i*-th position within the protein sequence, and *j* represents the 20 types of amino acids. Here, we calculated the absolute value of Δ*S* to reflect the magnitude of the impact of mutations on the compound-protein interaction.

After calculating the mutation activity change score, we evaluated whether a mutation at a specific position within the protein sequence plays an important role in compound-protein interactions. Therefore, we computed the average score of each position 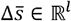 as

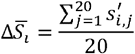

Furthermore, the value of 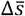 was normalized to between 0-1, and the relative activity change score Δ*Ri* ∈ ℝ^*l*^ was defined as

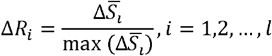

The relative activity change score Δ*R* can measure the contribution of each amino acid position for the compound-protein interaction.

### Experimental validation

#### Reagents and antibodies

Mouse antibody against GFP tag (catalog no. 66002-1-Ig), along with rabbit antibody targeting Beta Actin (catalog no. 66009-1-Ig), were pruchased from Proteintech. Secondary detection utilized HRP-conjugated Affinipure Goat Anti-Rabbit IgG (H+L) (catalog no. SA00001-2) and HRP-conjugated Affinipure Goat Anti-Mouse IgG (H+L) (catalog no. SA00001-1), also from Proteintech. All small molecules were sourced from MedChemExpress (MCE).

#### Cell culture and viral infection

The RAW264.7, and Hela cell lines were sourced from the American Type Culture Collection (ATCC).RAW264.7 were cultured in Dulbecco’s Modified Eagle’s Medium (DMEM) (EallBio, 03.1002C), while Hela were maintained in RPMI 1640 (EallBio, 03.4001C). All culture media were supplemented with 10% fetal bovine serum (FBS) and 100 U/ml penicillin-streptomycin for optimal cell growth and maintenance. Subsequently, the cells reaching 70-80% confluence were infected with Vesicular Stomatitis Virus (VSV) at a multiplicity of infection (MOI) of 0.1, Herpes Simplex Virus 1, F strain (HSV-1 F strain, MOI of 0.5), Encephalomyocarditis Virus (EMCV, MOI of 0.1), Mouse Hepatitis Virus, A59 strain (MHV-A59, MOI of 0.1), Influenza A Virus, PR8 strain (IAV-PR8, MOI of 0.1) and VSV expressing the green fluorescent protein (VSV-GFP, MOI of 0.1). After viral infection for 1 hour, the media were removed, and fresh medium was added to the cultures for further analysis.

#### RNA extraction, reverse transcription and real-time quantitative PCR

RNA extraction from cells following various infections was performed using TRIzol reagent (TIANGEN, A0123A01). The purified RNA was then reverse-transcribed into cDNA using HiScript II RT SuperMix (Vazyme, R223-01). Target gene expression was quantified using SYBR Green qMix (Vazyme, Q311) in a quantitative reverse transcription PCR (RT-qPCR) assay. The relative expression levels of target mRNAs were normalized to the housekeeping gene Actb. Detailed viral primer sequences used in this study are provided in **Supplementary Table 7**.

#### Total protein extraction and western blots analysis

Cells were lysed using RIPA Lysis Buffer (Strong) (MedChemExpress, HY-K1001), which was supplemented with protease inhibitors, including an EDTA-Free Protease Inhibitor Cocktail, Phosphatase Inhibitor Cocktail I (100× in DMSO), and Phosphatase Inhibitor Cocktail III (100× in DMSO). Clarified cell extracts (10 to 30 micrograms) were resolved on SDS-polyacrylamide gels and then transferred onto nitrocellulose membranes (Beyotime, FFN08). After blocking, the membranes were incubated with specific primary antibodies. The bound secondary antibodies were detected using the enhanced chemiluminescence (ECL) method (EallBio, 07.10009-50).

#### Flow Cytometry Analysis

After infection, cells were washed twice with phosphate-buffered saline (PBS) to remove residual virus, followed by trypsinization to generate a single-cell suspension. The cells were collected by centrifugation at 1600 rpm for 5 minutes, and the supernatant was discarded. The resulting cell pellet was resuspended in PBS and transferred to flow cytometry tubes for subsequent analysis. To evaluate the infection efficiency, the percentage of GFP-positive cells was quantified using a flow cytometer (BD Biosciences).

#### Fluorescence Assay

Cells were infected with VSV-GFP at an MOI of 0.1 for 12 hours. After treatment with small molecules, cells were visualized under a Nikon fluorescence microscope equipped with a 488 nm excitation filter and a 510 nm emission filter. Images were captured at a magnification of 10×.

#### Statistical analysis

In the present study, the Python package SciPy (https://pypi.org/project/scipy/) was used to conduct all the statistical analyses as indicated. Results are presented as the mean ± SEM. The p values < 0.05 were considered statistically significant (*), p values < 0.01, and p values < 0.001 were regarded as highly statistically significant (** and ***).

## Data availability

The antiviral-related bioactivity data based on cell assays were downloaded from the ChEMBL database (https://www.ebi.ac.uk/chembl/). The compound-protein binding affinity data were collected from the ChEMBL database and the BindingDB database (https://www.bindingdb.org/). The compound-protein complex structures were downloaded from the Protein Data Bank (PDB) database (https://www.rcsb.org/).

## Code availability

DeepAVC source code is available on GitHub (https://github.com/KangBoming/DeepAVC).

## Web server

We have developed a user-friendly web server that is freely accessible on the DeepAVC website (http://www.cuilab.cn/DeepAVC). The website was constructed on the basis of the packages of Python 3, Flask and Numpy.

## Acknowledgements

This study was supported by the grants from the National Natural Science Foundation of China [62025102, 32470924].

## Author information

These authors contributed equally: Boming Kang, Yang Zhao.

**Authors and Affiliations**

**Department of Biomedical Informatics, State Key Laboratory of Vascular Homeostasis and Remodeling, School of Basic Medical Sciences, Peking University, 38 Xueyuan Rd, Beijing, 100191, China**

Boming Kang, Rui Fan & Qinghua Cui

**Institute of Systems Biomedicine, Beijing Key Laboratory of Tumor Systems Biology, Department of Microbiology & Infectious Disease Center, NHC Key Laboratory of Medical Immunology, Peking University Health Science Center, Peking University, Beijing, China**.

Yang Zhao & Fuping You

**School of Sports Medicine, Wuhan Sports University, No. 461 Luoyu Rd. Wuchang District, Wuhan 430079, Hubei Province, China**

Qinghua Cui

**Contributions**

BK and YZ designed the study. BK and YZ performed the study. BK, YZ, RF, FY and QC wrote or edited the manuscript. QC and FY supervised the study.

**Corresponding author**

Correspondence to Fuping You & Qinghua Cui

## Figure Legends

**Extended Data Figure 1:**
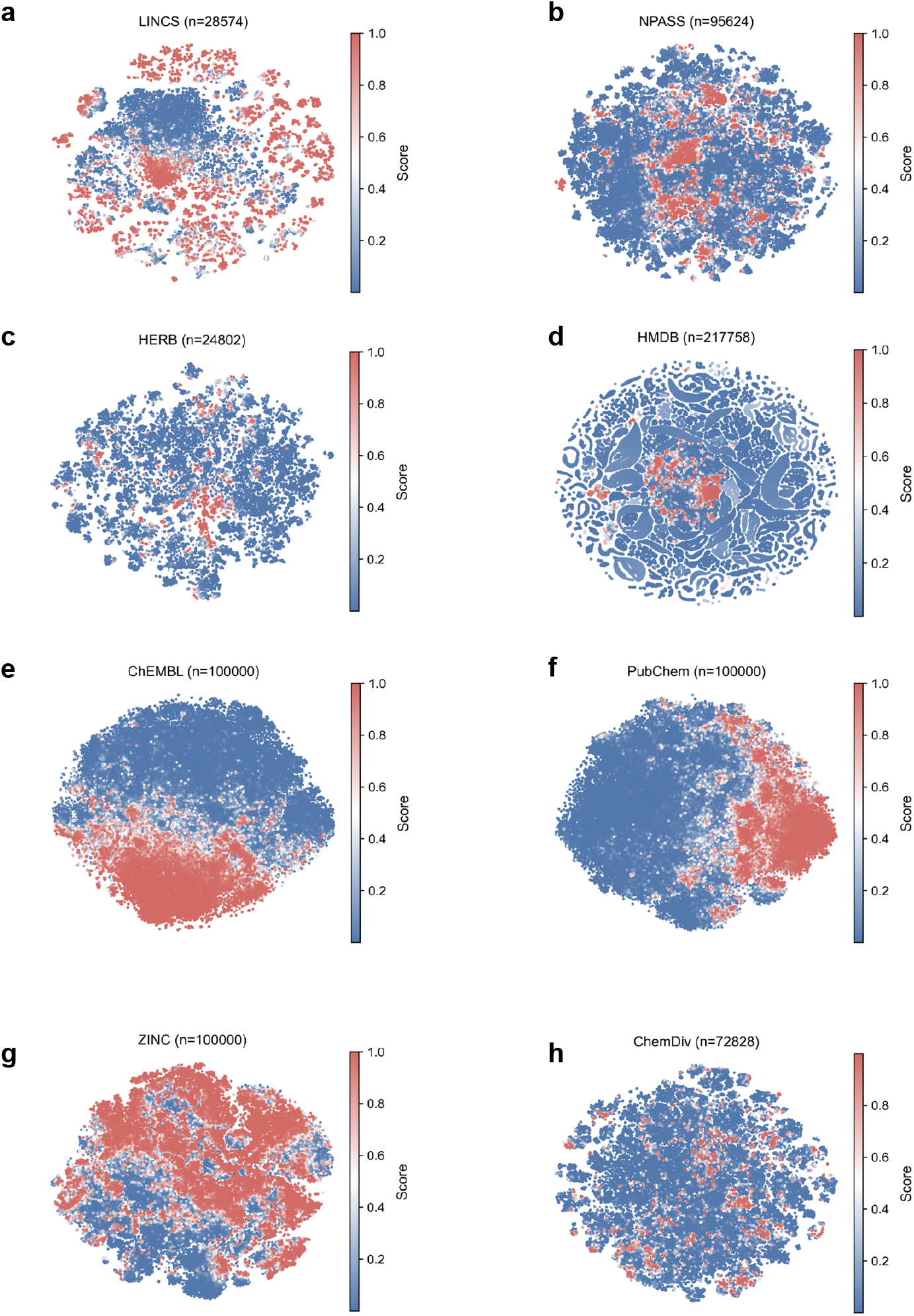
Virtual screening results of the remaining 8 large-scale chemical databases visualized by t-SNE. **(a)** The LINCS database. **(b)** The NPASS database. **(c)** The HERB database. **(d)** The HMDB database. **(e)** The ChEMBL database. **(f)** The PubChem database **(g)** The ZINC database **(h)** The ChemDiv database. The score refers to the antiviral score predicted by the DeepPAVC model. The ‘n’ denotes the number of compounds used for visualizing by t-SNE.

**Extended Data Figure 2:**
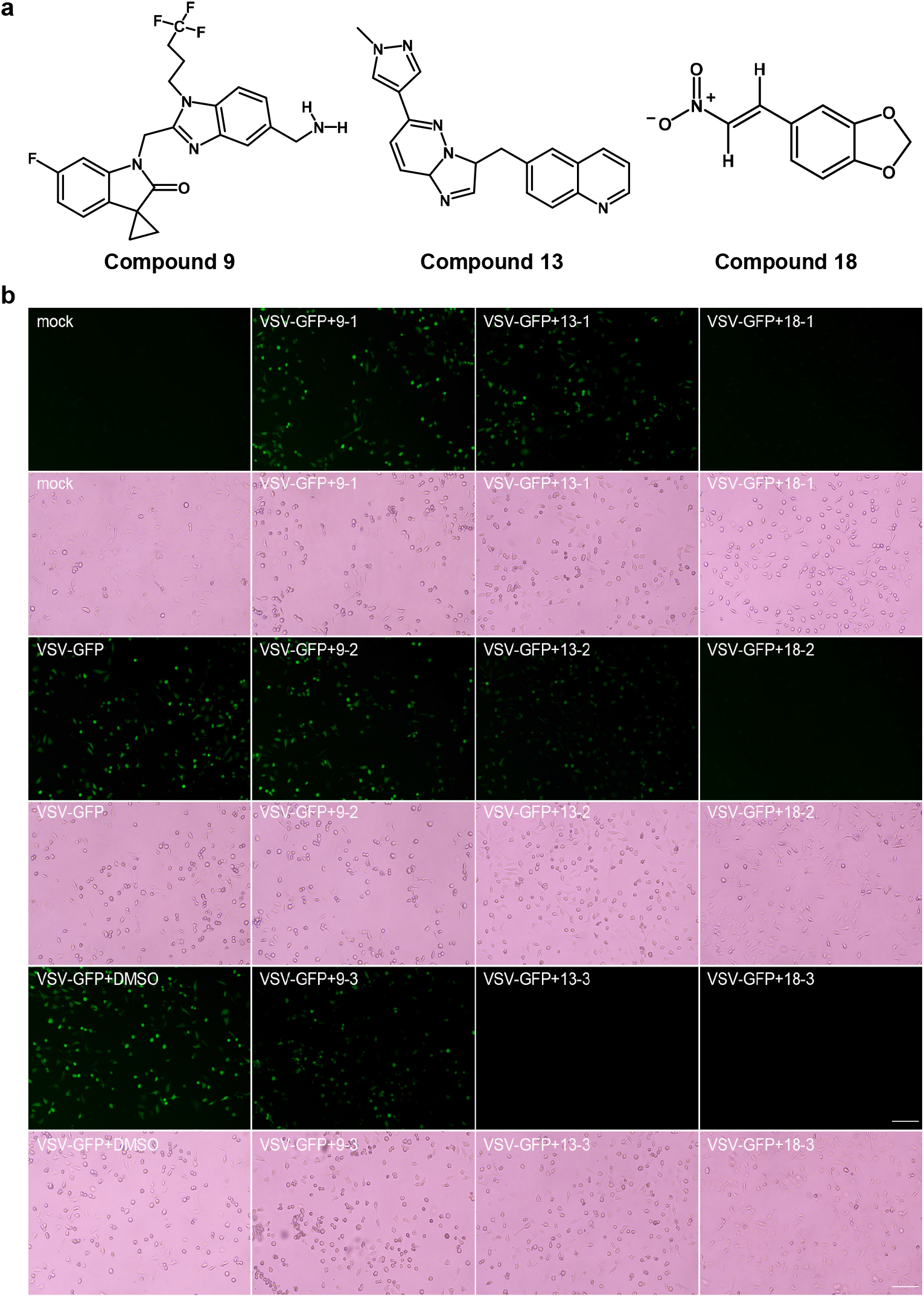
**(a)** The chemical structures of Compound 9, Compound 13 and Compound 18. **(b) Microscopic visualization of GFP fluorescence in HeLa cells treated with antiviral compounds**. HeLa cells were infected with VSV-GFP for 12 hours and treated with Compound 9, Compound 13, and Compound 18 at concentrations of 10 μM, 20 μM, and 40 μM, or DMSO as a control. Both bright-field and GFP fluorescence images were captured using a fluorescence microscope. The images demonstrate the impact of compound treatment on viral replication, with GFP fluorescence indicating the presence of the virus. Scale bar = 100 μm.

